# Functional Near Infrared Spectroscopy (fNIRS) based Odor Detection and Classification using Functional Data Analysis

**DOI:** 10.1101/2021.02.13.431051

**Authors:** Faezeh Moradi, Shima T. Moein, Issa Zakeri, Kambiz Pourrezaei

## Abstract

An objective approach for odor detection is to analyze the brain activity using imaging techniques during the odor stimulation. In this study, Functional Near Infrared Spectroscopy (fNIRS) is used to record hemodynamic response from the frontal region of the brain by using a 4-channel fNIRS system. The fNIRs data is collected during the odor detection task in which the subjects were asked to press a button when they detect the given odor. Functional Data Analysis (FDA) was applied on fNIRs data to convert discrete measured samples of data to continuous smooth curves. The FDA method enables us to use the bases coefficients of fNIRS smoothed curves for features that represent the shape of the raw fNIRS signal. With the learning algorithm that we proposed, these features were used to train the support vector machine classifier. We evaluated the odor detection problem, in two binary classification cases: odorant *vs*. non-odorant and odorant *vs*. fingertapping. The model achieved a classification accuracy of 94.12% and 97.06% over the stimulus condition in the two cases, respectively. Moreover to find the actual predictors we used the extracted defined features (slope, standard deviation, and delta) to train our classifier. We achieved an average accuracy of 91.18 % on classifying odorant *vs*. non-odorant and an accuracy of 94.12% for odorant *vs*. fingertapping on the stimulus condition. The results determined that fNIRs signals of odorant and non-odorant are distinguishable without being affected by the motor activity during the experiment.

These findings suggest that fNIRs measurement on the forehead could be potentially used for objective and comparably inexpensive assessment of odor detection in cases that the subjective report is unreliable.

## 1. Introduction

Among medical brain imaging techniques, functional near infrared spectroscopy (fNIRS)^1,2,3^, with its excellent sensitivity to hemoglobin and lower sensitivity to motion artifact, is a non-invasive, inexpensive, portable technique that makes it more clinic friendly for applications.

To the best of our knowledge, Loreman and Pourrezaei were the first group that have used fNIR to study human hemodynamic response to odorant. In the olfactory stimulation, they found brain activity, in probes located on the frontal and temporal region.^4^

Due to the hemodynamic response of odorant on the frontal region measured by fNIRS and to its ease of measurement in that area, in this study, we use fNIRS measurement for odor assessment.

The hemodynamic responses measured by fNIRS are naturally continuous physiological processes that are observed in discrete time with the presence of observational noise. The recorded sampling points are likely to be correlated. Therefore, it is desirable to consider the discrete measures samples over time intervals as a continuous smooth function. Function Data Analysis (FDA) is a time series technique that its approach is to construct a smooth functional form from a set of discrete measured values that are distributed over a continuum which is often time.^5^ Utilizing FDA approach on fNIRS data was first introduced in Barati et al^6^. In another research study, Pourshoghi et al.^7^ proposed an application of FDA on fNIRS signal for the purpose of pain assessment.

In this study, we assumed that the hemodynamic changes of odorant and non-odorant signals are different and due to that we construct a classifier for the odor detection problem.

To test this hypothesis we use FDA to construct features from the data described at Sec. 2.1 and with the learning algorithm proposed in Sec. 2.4 we will construct a classifier for the odor detection problem.

## 2. Materials and Methods

### 2.1 Participants

Seventeen healthy individuals aged between 20 to 40 years old (10 female) participated in this study. All subjects performed the University of Pennsylvania Smell Identification Test (UPSIT) and were identified as normosmic. The protocol of the experiment was approved by the institutional ethics committee of Institute for Research in Fundamental Sciences (IPM).

### 2.2 Experimental Protocol

The experiment consisted of three blocks each with 7 interleaved presentations of stimulus odorant bottles (containing 0.5 mL of strawberry or garlic) or non-odorant bottles (odourless water). In each trial, the subject had to press a button as soon as the odorant was detected. At the end of each block after the presentation of odorant and non-odorant bottles, the subject was cued once with a light tapping on hand to make up to 10 finger tappings with intervals decided by the subject. After the end of each block the subject could take rest up to 15 minutes.

We consider three different conditions for each trial: (1) a duration of 5 seconds before the subject understands the odor, known as prestimulus; (2) the stimulus condition that starts from the time the subject understands the odor to 15 seconds after and (3) the 5 seconds recorded after the stimulus condition, known as poststimulus.

### 2.3 Measurements

The collected data^8^ were acquired using a custom-made 4-channel fNIRS system at the olfactory laboratory at Institute for Research in Fundamental Sciences (IPM). The probe with two near channels (ch1 and ch3) and two far channels (ch2 and ch4) was positioned on the forehead of the participant and with a sampling rate of 7.5 Hz. The probe consisted of two light-emitting diodes (LED) sources for infrared wavelengths located on the right and left side. Two infrared detectors were placed at 1.3 cm from the LED to make near channels to investigate the hemodynamic response and one infrared detector was located at 2.8 cm from LED to make far channels to measure the hemodynamic responses towards the midline. Each LED emitted two wavelengths of 730 and 850 nm.

### 2.4 fNIRS Data Processing

The optical density time series was first band-pass filtered with a cut-off frequency of 0.01-0.5 to remove the instrumental and physiological noise. The oxyhemoglobin (Hbo2) and deoxyhemoglobin (HB) concentration changes were obtained using the modified Beer-Lambert law.^9^

The following methods are applied on the average over trials of Hb and Hbo2 for each subject.

### 2.5 Functional Data Analysis (FDA)

In functional data analysis, the approach is to construct a smooth functional form from a set of discrete measured values *y*_*i*1_,.., *y_in_* that are distributed over continuum (e.g. time). Owing to the presence of measurement error in the raw discrete data we assume the following mathematical model for one recorded sample

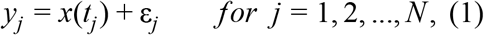

Here, *y_j_* is the measured sample at time *t_j_*, *x*(*t_j_*) value of smooth function that we want to determine and ε_*i*_ is the measurement error of the sample at time *t_j_*.

At this point FDA tries to find a smooth function that best describes the discrete data by minimizing the measurement error under a defined smoothness condition.^10^ In this study we use the spline smoothing method to take control over the level of smoothness. Spline smoothing is the most popular and powerful approach with well-defined properties.^11,12^

With the sum of squares criterion

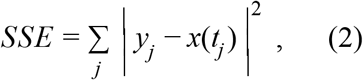

the closeness of the fitted curve to raw data is measured; However, we don’t want the curve to interpolate the data. Spline smoothing controls the degree of smoothness with the roughness penalty to achieve an improvement in the squared error cost function.

A popular way to quantify the roughness of a function is the square of the second derivative [*D*^2^*x*(*t*)] (*D*^2^ is an operator for second derivative) of the function at time *t*, is often called its curvature at *t*.^5^ The measure of function’s roughness is the integrated curvature criterion

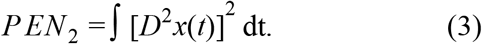

Therefore, by adding the roughness penalty to Eqs. (2) the new penalized sum of squares criterion will change to

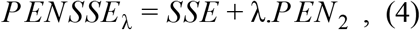

where *λ* is the smoothing parameter that measures the balance between the closeness to the raw data and the variability of the fitted curve to the data.

The larger value of *λ*, the more the spline fit is diminished toward the linear regression estimation of data. Therefore, it’s important to estimate an appropriate smoothing parameter. The smoothing parameter estimation is done after determining the number of knots in the functional form of data (more detail in the section 2.2.1).

The smoothing parameter can be chosen subjectively by visual judgment and prior knowledge of the process of generating the data. An objective, data-driven method is also developed by using the generalized cross validation (GCV) measure.^13^

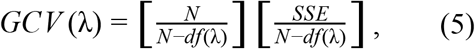

where N is the number of observations, SSE is the mean square error, and *df*(*λ*) is the trace of the smoothing matrix.^14^

#### 2.5.1 Building a continuous and smooth function from discrete data

A Basis function system is used to represent the functional form of the raw discrete data. A basis function system is a set of known functions ϕ_*k*_ that are mathematically independent of each other. This basis function system has the property of approximating arbitrarily well any function by taking a weighted sum or linear combination of a sufficiently large number K of these functions.^5^

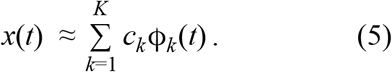

The parameters *c*_1_,.., *c_k_* are coefficients and ϕ = (ϕ_1_, …, ϕ_*K*_) is the vector of length k containing the basis functions.

There are several basis functions available like Fourier series, B-spline, polynomial and exponential. The choice of basis function depends on the behaviour of our raw data. For a physiological signal like hemodynamic response B-spline basis system is considered as the best option.^5^

B-spline basis system is constructed from spline basis function. A spline function is a piecewise polynomial function developed between knots. The first step in defining a spline is to divide the interval over which a function is to be approximated in each subinterval. The time points in which these subintervals join are called knots. Over each interval, a polynomial of specified degree *m* is defined. The order (one more than its degree; *m* + 1) of a polynomial is the number of constants required to define it. In other words, it is the number of bases coefficients that define the function.

To sum up, a spline system is determined by two things: the order of polynomials *m*, and the knot sequence (number of knots and the location of knots).

The goal for any algorithm in selecting the number of knots (K) is to make certain that K is sufficiently large to the data and not so large that computation time is unnecessarily large. We determine the number of knots according to the “myopic” algorithm. The “myopic” algorithm stops when no improvement in the generalized cross validation statistic (GCV) is noticed with the last increase in the number of knots.^15^

All of the functional data analysis implementations are conducted in MATLAB by using the FDA software package in MATLAB.^14^

### 2.6 Feature selection method

Feature selection methods can significantly improve classification accuracy by selecting the most informative features. In this study, we used a recursive feature elimination method by training on the support vector machine classification model (RFE_SVM).

RFE_SVM removes irrelevant features based on backward elimination procedure. The output of this feature selection method is the rank list with features that are ordered based on their predictive performance.^16^

Our implementation of RFE_SVM is described in the following steps (FeatureSelectionProcedure):

1. SVM model selection (Parameter tuning)
2. Train the SVM model.
3. Calculate weight vector of the SVM model and define the ranking criteria.
4. Remove the variable or feature with the lowest ranking criteria value.
5. go to step 3 until the stopping criteria.

### 2.7 Learning algorithm

The learned classifier can predict the corresponding class label of the new samples by only giving the new samples features (predictors). The learned classifier does this prediction by approximating a mapping function from the features determined from the feature selection method (predictors) to its corresponding class label (target).

In this study, our focus is on a binary classification problem in which the class or target is limited to two values.

The support vector machine (SVM) classifier computes a separating hyperplane that discriminates by maximizing the separating margins between the hyperplane and the nearest data point in each class. For a given training set we have a pair of (*X_i_,y_i_*), *X_i_* ∈ *R^n^, y_i_* ∈ {1,-1}, *i* =1,.., *l*, which means each predictor is tagged to its class label. The hyperplane is found by solving the following optimization problem:^17^

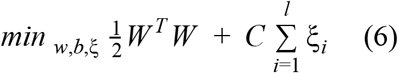

Subject to *y_i_*(*W^T^*ϕ(*X_i_*) + *b*) ≥ 1 − *ξ_i_*, *ξ_i_* ≥ 0 where ϕ() is the function that transforms the training data *X_i_* into higher dimensional space, *ξ_i_* is the slack variable to manage no-separable cases, *W* is the weight matrix, b is bias term and *C* is the regularization parameter that determines the tradeoff between margin width and training error.^18^ In the cases that classes are non-linearly separable (ϕ(*x*) ≠ *x*), SVM uses a kernel function to calculate a non-linear separating boundary. In our experiment, we used a radial basis function (RBF) kernel defined as

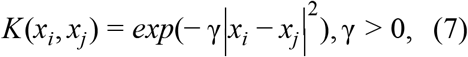

which leads to a support vector machine with RBF kernel (SVM_RBF) classifier.

The learning algorithm, which we apply on our binary classification problem, has the following paradigm [^18,19^, Section 7.10]:

1. SVM_RBF model selection: Select the best values for kernel parameter (γ) and regularization parameter (*C*) in the SVM (RBF kernel) classifier.
2. Divide the samples into separate groups as follows: each time remove a subject with all its instances from the training dataset and use it for the testing dataset.
3. Iterate over the groups: k=1,..,K (K is the number of subjects)
4. Apply the “FeatureSelectionProcedure” (mentioned in sec. 2.3) to find a subset of good predictors.5. Using just this subset of predictors, to train the classifier (the selected model in step 1).
5. Use the learned classifier model to predict the class labels for the samples of the removed subject.

The parameter tuning is the process in which you search for the best parameter using the train data so that the classifier can accurately predict unknown data (testing data). Various pairs of (γ, *c*) values among the ranges *c* = [2^−5^,2^−3^,.., 2^15^], γ = [2^−15^,2^−13^,.., 2^3^] are tried and the one with the best cross-validation accuracy is picked. ^20^

We used leave one out cross validation in the proposed learning algorithm that has the effect to generalize the results to new dataset.

The machine learning algorithms are all implemented in R by using the svm package e1071.

## 3. Results

### 3.1 Functional data analysis results

Functional data analysis is used to convert Hb and Hbo2 olfactory time-series data to their functional form. With this conversion the 188 raw discrete data points will be reduced to the number of bases coefficients of the functional form. We use the coefficients as a representation for the olfactory raw data for our future classification study, this allows us to study the shape of the fitted curve.

We found a B-spline basis system with order 3, sufficient to fit the raw data.

We have determined the knots to be equally spaced on the interval and the number of knots are evaluated from 5 to 30 with the step size 1 according to the “myopic” algorithm.

As mentioned in section (2.1) the raw data is splitted into three parts, so the number of knots is determined for each part separately. We selected 8 knots for the prestimulus and poststimulus condition and 23 knots for the stimulus condition.

Fig. 1 also verifies the selected number of knots for the stimulus condition by evaluating the changes of knots over the classification accuracy of the odorant and non-odorant classification problem.(more detail of classification is described in sec. 3.3).

**Fig. 1.**
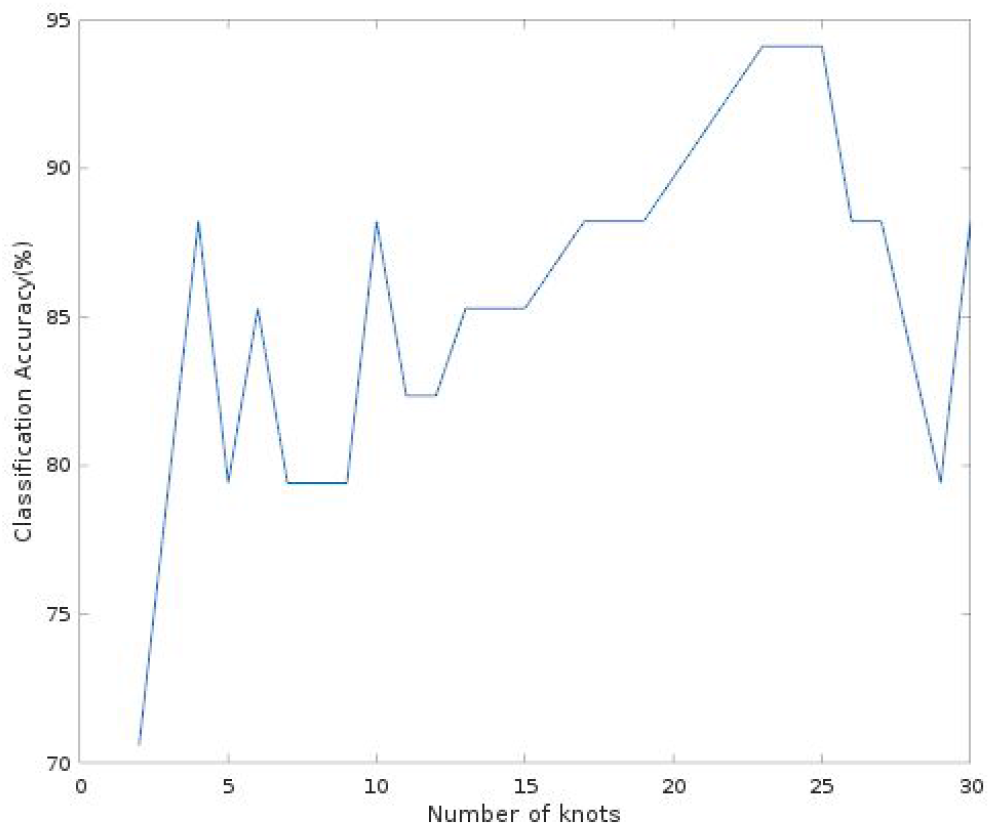
The effect of changing the number of knots on odorant vs. non-odorant classification accuracy on the stimulus condition

### 3.2 Feature construction

In this study, we use the bases coefficients of the smooth fitted curve of Hb and Hbo2 data, to construct the feature set.

As mentioned in Sec. 3.1, B-spline of order 3 with 23 numbers of knots is used for the FDA setting on the raw data (stimulus condition). The number of parameters required to represent this B-spline system is the order plus number of interior knots^5^ (total number of knots minus two), which gives 24 bases function in this case.The data is measured over 4 channels and on each channel two signals Hb and Hbo2 are extracted. Each trial will have a coefficient matrix of size 192 over the stimulus condition. Therefore the feature space dimensionality of the data is measured as,

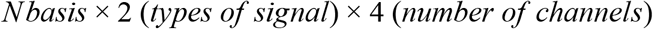

where Nbasis is the number of bases function. Fig. 2 illustrates how the feature set is constructed.

**Fig. 2.**
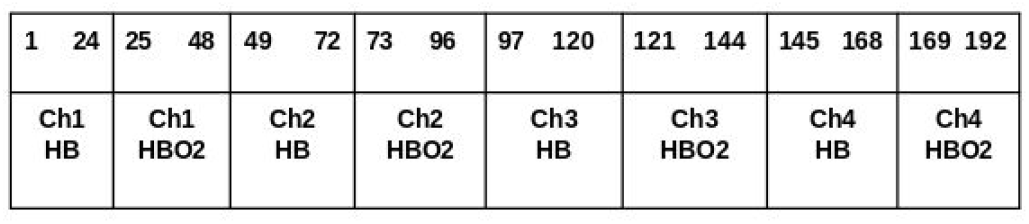
Feature set for stimulus part of Hb and Hbo2: A concatenation of the functional form bases coefficients over signals (Hb and Hbo2) and channels (channel 1, channel 2, channel 3, and channel 4)

The same is applied to prestimulus and poststimulus conditions.

### 3.3 Feature extraction

In this part, we define some features that represent the shape of the Hb and Hbo2 signals, these features are explained as below,

1. Δ: changes of the signal during the stimulus condition.
2. Slope: is the slope of the stimulus condition. Two slopes are defined, one from 0 to 5 seconds and the other from 5 to 15 seconds.
3. SD: is defined as the ratio of the standard deviation of the poststimulus condition over the standard deviation of the prestimulus condition.

The above defined features are calculated for Hbo2, Hb, and the ratio of Hbo2 and Hb signals. With this in mind, we have 12 features for each channel, which will be 48 features for all four channels.

### 3.4 Classification results on FDA coefficients

In this section, we will present the classification performance of the proposed method in Sec. 2.4 by using the coefficient of the smooth fitted curve of Hb and Hbo2 data collected by fNIRS.

Results in Table 1 show that the maximum classification accuracy is achieved over the stimulus condition, from the time which the participant is presented with the stimulus bottle to 15 seconds after.

**Table 1.** Results of classification accuracy and number of bases function for each condition.

|  | Odorant vs. Non-odorant |  | Odorant vs. finger tapping |  |
| --- | --- | --- | --- | --- |
|  | Accuracy (%) | Nbasis | Accuracy (%) | Nbasis |
| Prestimulus | 73.53 | 9 | 85.29 | 9 |
| Stimulus | 94.12 | 24 | 97.06 | 24 |
| Poststimulus | 85.29 | 9 | 64.71 | 9 |
| Whole episode<br>(Prestimulus,<br>Stimulus and<br>Poststimulus) | 91.18 | 33 | 85.29 | 33 |

The accuracy of 94.12% is obtained for classifying odorant and non-odorant on the stimulus condition. This indicates that the hemodynamic changes of odorant signals are different from non-odorant signals. As explained in Sec. 2.2 the experiment includes the finger tapping action that might affect the recorded fNIRS odorant signals. As a result, a SVM_RBF classifier is constructed to evaluate how distinguishable odorant and finger tapping signals are, by using the coefficient features. From Table 1 the accuracy of 97.06% is achieved for classifying odorant and finger tapping signals. Therefore, we conclude the difference between the odorant and non-odorant is not affected by motor activity while the subject is smelling.

The bar plot in Fig. 3 shows the classification accuracy of using only near channels (channel 1 and channel 3) that measures skin responses, using far channels (channel 2 and channel 4) that measures mostly cortex and a bit skin response and both considering far and near channels.

**Fig. 3.**
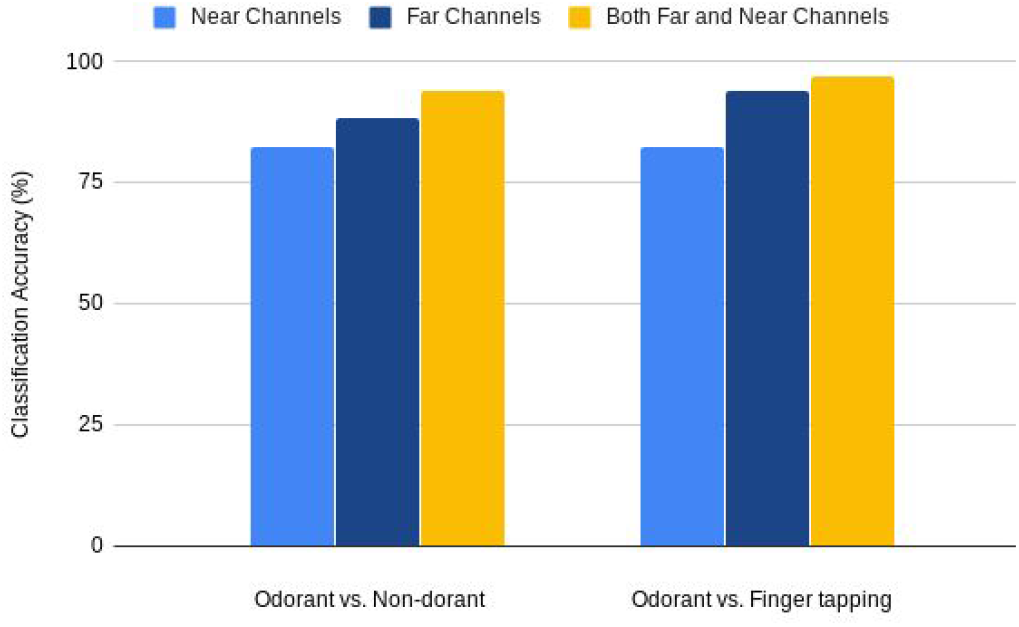
Classification accuracy by using only near channels, only far channels and both far and near channels are compared on three classification problems : odorant *vs*. non-odorant and odorant *vs*. finger tapping.

The results show that using both far and near channels give a higher classification accuracy. This is true for all two observations: odorant *vs*. non-odorantand and odorant *vs*. finger tapping.

### 3.5 Classification results on extracted features

In order to obtain our actual predictors, we used the extracted features defined in Sec. 3.2. On the new feature set, the SVM_RFE feature selection method is applied to determine the most informative features. With this new representation for Hb and Hbo2 we built a SVM_RBF classier model by using the extracted features.

We believe that the extracted features based on the smoothed curves shape can acquire a high accuracy to the same result achieved by using coefficient features. In the case of classifying odorant and non-odorant, as expected we achieved an average accuracy of 91.18% on the stimulus condition. And an accuracy of 94.12% for classifying odorant and finger tapping, This indicates odorant and non-odorant signals are not affected by finger tapping.

## 4. Conclusion

Functional near infrared spectroscopy (fNIRS) is a powerful tool for measuring hemodynamic changes of living brain tissues.

In this study, we have used fNIRS measurement on the frontal region for odor assessment. Beside the natural continuous form of fNIRs data, the analysis of fNIRS data is limited to discrete time series methods.

In order to study the functional form of fNIRs signal, with the use of functional data analysis (FDA) we converted the discrete fNIRS data to its continuous time functional form. This helped us to extract features that represent the raw fNIRS signal from its smoothed curve and with the proposed learning algorithm explained in Sec. 2.4 we trained the SVM_RBF classifier to achieve a model for the odor detection problem.From the results presented in section 3.3 we could conclude that the odorant and non-odorant signals are distinguishable according to the shape of the fitted curve of fNIRS data with a classification accuracy of 94.12%, over the coefficient selected features.

Results on odorant and fingertapping indicate that the odor signal is not affected by the motor activity during the experiment. The classification accuracy results are 97.06% and 94.12% respectively over coefficient selected features and extracted selected features.

## Acknowledgments

I would like to thank Dr. Shima Moein for her key role in experiment design and collection of the data presented in this paper. I would like to express my gratitude to Prof. Issa Zakeri for all valuable comments and guidance throughout this research.

